# Uncoupling protein 1-driven Cre (*Ucp1-Cre*) is expressed in the epithelial cells of mammary glands and various non-adipose tissues

**DOI:** 10.1101/2023.10.19.563175

**Authors:** Kyungchan Kim, Jamie Wann, Hyeong-Geug Kim, Jisun So, Evan D. Rosen, Hyun Cheol Roh

**Author notes:** Corresponding author: Address: 635 Barnhill Drive, MS1021G, Indianapolis, IN 46202, USA.

## Abstract

**Objective:** Uncoupling protein 1 (UCP1), a mitochondrial protein responsible for nonshivering thermogenesis in adipose tissue, serves as a distinct marker for thermogenic brown and beige adipocytes. *Ucp1-Cre* mice are thus widely used to genetically manipulate these thermogenic adipocytes. However, evidence suggests that UCP1 may also be expressed in non-adipocyte cell types. In this study, we investigated the presence of UCP1 expression in different mouse tissues that have not been previously reported.

**Methods:** We employed *Ucp1-Cre* mice crossed with Cre-inducible transgenic reporter Nuclear tagging and Translating Ribosome Affinity Purification (NuTRAP) mice, to investigate *Ucp1*-*Cre* expression in various tissues of adult female mice and developing embryos. Tamoxifen-inducible *Ucp1-CreERT2* mice crossed with NuTRAP mice were used to assess active UCP1 expression. Immunostaining, RNA analysis, and single-cell/nucleus RNA-seq (sc/snRNA-seq) data analysis were performed to determine the expression of endogenous UCP1 and *Ucp1-Cre*-driven reporter expression. We also investigated the impact of UCP1 deficiency on mammary gland development and function using *Ucp1*-knockout (KO) mice.

**Results:** *Ucp1-Cre* expression was observed in the mammary glands within the inguinal white adipose tissue of female *Ucp1-Cre*; NuTRAP mice. However, endogenous *Ucp1* was not actively expressed as *Ucp1-CreERT2* failed to induce the reporter expression in the mammary glands. *Ucp1-Cre* was activated during embryonic development in various tissues, including mammary glands, as well as in the brain, kidneys, eyes, and ears, specifically in epithelial cells in these organs. While sc/snRNA-seq data suggest potential expression of UCP1 in mammary epithelial cells in adult mice and humans, *Ucp1*-KO female mice displayed normal mammary gland development and function.

**Conclusions:** Our findings reveal widespread *Ucp1-Cre* expression in various non-adipose tissue types, starting during early development. These results highlight the importance of exercising caution when interpreting data and devising experiments involving *Ucp1-Cre* mice.

## 1. INTRODUCTION

Adipose tissues play a crucial role in energy balance and metabolic homeostasis. These tissues not only store energy in the form of lipids but also function as an endocrine organ by secreting various hormones and cytokines that regulate energy metabolism and insulin sensitivity [1].

Adipose tissue is broadly classified into two types, white adipose tissue (WAT) and brown adipose tissue (BAT). WAT primarily functions in the role of energy storage, whereas BAT dissipates energy through nonshivering thermogenesis for thermal regulation. While these two adipose tissue types are located in anatomically distinct locations, a third type of adipocyte—the thermogenic beige adipocyte—has recently been identified to emerge within WAT upon cold exposure [2]. These thermogenic brown and beige adipocytes have gained significant attention due to their potential to counteract obesity and related metabolic disorders [3].

The thermogenic function of brown and beige adipocytes is primarily contributed by mitochondrial uncoupling protein 1 (UCP1) [4]. UCP1 uses the proton gradient across the mitochondrial inner membrane to generate heat by uncoupling oxidative phosphorylation from ATP production. While recent studies have revealed UCP1-independent thermogenic mechanisms [5,6], UCP1 expression remains a hallmark of thermogenic adipocytes. The study of these thermogenic adipocytes has been greatly facilitated by the development of the *Ucp1-Cre* transgenic mouse line [7], allowing for genetic manipulation and labeling of UCP1-expressing thermogenic adipocytes *in vivo*, significantly advancing our understanding of adipose biology.

Accumulating studies have documented the expression of UCP1 in various non-adipose tissues, such as the kidneys, adrenal glands, skeletal muscle, heart, thymus, brain, and gastrointestinal tract [8–15]. Although the extent and functional implications of this expression are still under investigation, the presence of UCP1 in these non-adipose tissues suggests potential roles beyond thermogenesis, including metabolic regulation and protection against oxidative stress [9]. Obviously, the expression of UCP1 in non-adipose tissues raises concerns regarding the tissue specificity of *Ucp1-Cre* for generating mouse models with brown/beige adipocyte-specific genetic perturbations. Indeed, Claflin et al. observed the expression of *Ucp1-Cre* not only in adipose tissues but also in other tissues, such as the kidneys, adrenal glands, and hypothalamus [8]. This suggests that there may be potential off-target effects on non-adipose tissues when using *Ucp1-Cre* mice for specific targeting of brown/beige adipocytes. In our study, we report the expression of *Ucp1-Cre* in myoepithelial and ductal epithelial cells of the mammary gland located within the inguinal WAT (iWAT) of female mice. The expression of *Ucp1-Cre* in the mammary glands occurs during embryonic development. *Ucp1-Cre* is also found in epithelial cells in various non-adipose tissues in developing embryos, including the eyes, ears, whisker follicles, and brain, although there is no active endogenous UCP1 expression in adult tissues. Single-cell and nucleus RNA sequencing (sc/snRNA-seq) data suggest the possibility of transient UCP1 expression in mammary glands postnatally in mice and humans. Nevertheless, *Ucp1*-knockout (KO) mouse studies suggest that UCP1 is not necessary for the development or function of the mammary glands. Our study highlights that it is crucial to consider the broad expression pattern of *Ucp1-Cre* in studies that employ mouse models utilizing this genetic tool.

## 2. MATERIAL AND METHODS

### 2.1. Animals

All animal experiments were performed according to procedures approved by the Beth Israel Deaconess Medical Center (BIDMC) Institutional Animal Care and Use Committee and the Indiana University School of Medicine (IUSM) Institutional Animal Care and Use Committee. To label UCP1-expressing cells, NuTRAP (Nuclear tagging and Translating Ribosome Affinity Purification) mice (Jackson Laboratory, 029899) [16] were crossed with *Ucp1-Cre* mice (Jackson Laboratory, 024670) [7] and *Ucp1-CreERT2* mice [17], respectively. Mice were housed under a 12-hour light/12-hour dark cycle with ad libitum access to a standard chow diet and water at 22°C for standard room temperature housing. To evaluate the Cre recombinase activity in response to cold and thermoneutrality, mice were housed in 4°C and 30°C incubators for 1 week and 4 weeks, respectively. *Ucp1*-KO mice (Jackson Laboratory, 003124) were used for lactation assays.

### 2.2. Tamoxifen injection

Tamoxifen (Sigma, T5648) was dissolved in sunflower seed oil (Sigma, S5007) at a concentration of 20 mg/mL by incubating at 37°C overnight with agitation. The mice were then intraperitoneally injected with tamoxifen at a dosage of 100 mg/kg for three consecutive days. After a washout period of 3 days following the last injection, the mice were euthanized to collect tissue samples.

### 2.3. Lactation assay

*Ucp1*-KO or heterozygous female mice were mated with wild-type C57BL/6 male mice, and then body weight measurements of their offspring were taken during the 2-week lactation period. After weaning, female breeders were sacrificed for histological analysis of iWAT.

### 2.4. Immunohistochemistry

Dissected tissues and embryos were fixed in 10% formalin for 1-2 days at 4°C, washed in PBS, and processed for paraffin-embedded sections by the BIDMC or IUSM histology core. For immunohistochemical staining, sections were deparaffinized, treated with sodium citrate for antigen retrieval, and then with hydrogen peroxide to inactivate endogenous peroxidase. The sections were subsequently blocked with donkey serum and incubated with rabbit anti-GFP antibody (1:500; Abcam, ab290) or rabbit anti-UCP1 antibody (1:500; Abcam, ab10983) at 4°C overnight. After extensive washing in PBS with 0.05% Tween-20, the sections were incubated with an anti-rabbit secondary antibody, enhanced, developed in diaminobenzidine, and counter-stained with hematoxylin.

### 2.5. Immunofluorescence staining

Dissected tissues were fixed in 4% paraformaldehyde in PBS for 1 day at 4°C and then immersed in 30% sucrose in PBS for 2 days. To prepare frozen sections, the tissues were embedded in an optimal cutting temperature compound (Sakura Finetek, 4583) and frozen on dry ice. Serial sections (40 μm) were cut on a cryostat (Leica, CM1950), mounted on adhesive glass slides (Matsunami, SUMGP11), and air dried for 2 hours at room temperature. After extensive rinsing in PBS, the sections were incubated in a mixture of goat anti-GFP (1:300; Novus, nb100-1678) and rabbit anti-UCP1 (1:300; Abcam, ab10983) primary antibodies diluted in PBS with 5% donkey serum and 0.1% Triton X-100 at 4°C overnight. After extensive rinsing in PBS with 0.05% Tween-20, Alexa488-conjugated donkey anti-goat (1:300; Invitrogen, A11055) and Alexa568-conjugated donkey anti-rabbit (1:300; Invitrogen, A10042) secondary antibodies were applied for 2 hours at room temperature. The sections were subsequently incubated for 10 minutes with 1 μg/mL Hoechst 33258 (Invitrogen, Thermo Fisher Scientific) in PBS for nuclear staining, then mounted on glass slides, and coverslipped with VECTASHIELD Mounting Medium (Vector Laboratories). Sections were visualized under Lecia TCS SP8 confocal laser-scanning microscope.

### 2.6. RNA extraction and qRT-PCR

TRIzol reagent (Ambion, Thermo Fisher Scientific) and chloroform were used to extract total RNA from each tissue, according to the manufacturer’s protocol. cDNA synthesis and qRT-PCR analysis were performed as we have described previously [18]. The primer sequences used for qRT-PCR are as follows: *Ucp1*: Forward-ACTGCCACACCTCCAGTCATT, Reverse-CTTTGCCTCACTCAGGATTGG, *GFP*: Forward-GCGGCGGTCACGAACTC, Reverse-AGCAAAGACCCCAAGAGAA, *36B4*: Forward-GAGGAATCAGATGAGGATATGGGA, Reverse-AAGCAGGCTGACTTGGTTGC.

### 2.7. Single cell/nucleus RNA-seq (sn/scRNA-seq) analysis

The snRNA-seq data from mouse adipose tissue [19] were accessed and analyzed using the Single Cell Portal provided by the Broad Institute (https://singlecell.broadinstitute.org/single_cell). The human scRNA-seq data [20] were accessed through the Human Protein Atlas (https://www.proteinatlas.org). The UCP1 expression data were retrieved directly from the databases without any additional processing.

### 2.8. Statistics

Two-tailed unpaired Student’s *t* test was carried out using GraphPad Prism 8. Data are presented as mean ± SEM and significance was determined at *P* < 0.05.

## 3. RESULTS

### 3.1. *Ucp1-Cre* is expressed in the mammary glands of female mice

We previously developed a transgenic reporter mouse called NuTRAP. This mouse harbors a transgene cassette at the *Rosa26* locus that produces GFP-tagged ribosomes and mCherry-tagged nuclei. The cassette is preceded by a loxP-flanked stop sequence, enabling transgene expression in a Cre recombinase-dependent manner [16]. This mouse is useful for conducting transcriptional and epigenomic analyses in a cell type-specific manner [21]. As Cre-mediated recombination removes the stop sequence and subsequently leads to permanent labeling of the cells, it can be utilized for lineage tracing studies by tracking GFP/mCherry expression. To investigate UCP1-expressing beige adipocytes, we generated NuTRAP crossed with *Ucp1-Cre* mice and exposed these mice to cold conditions for 1 week. Following immunohistochemical staining of GFP in the male mice subjected to cold temperatures, we observed specific GFP expression in dense multilocular beige adipocytes within the iWAT depot (**Figure 1A**), which is consistent with our previous study [22]. Interestingly, female *Ucp1-Cre*; NuTRAP mice exhibited GFP expression not only in beige adipocytes but also in cell types within the mammary gland, including myoepithelial and luminal epithelial cells (**Figure 1B**). The abundance of GFP-expressing beige adipocytes was significantly changed in response to different housing temperatures: highest in the cold, intermediate at room temperature, and lowest under thermoneutral conditions (**Figure 1C**). However, GFP-expressing cells in the mammary gland did not exhibit a temperature-dependent response (**Figure 1C**). When we conducted UCP1 immunohistochemical staining on consecutive iWAT sections from the cold-exposed female *Ucp1-Cre*; NuTRAP mice, we found UCP1 staining in both beige adipocytes and mammary gland cells, similar to the GFP staining (**Figure S1A**). Nonetheless, there was high background staining (**Figure S1A**), which raised concerns about the specificity of the UCP1 antibody. To address this issue, we performed UCP1 immunohistochemical staining on iWAT from female *Ucp1*-KO mice. As previously reported [23,24], we observed a higher number of multilocular beige adipocytes in the iWAT of *Ucp1*-KO mice compared to the control *Ucp1* heterozygous mice (**Figure S1B**). These beige adipocytes in *Ucp1*-KO mice were negative for UCP1 expression. However, UCP1 immunoreactivity was still evident in the mammary glands of both *Ucp1*-KO and heterozygous mice (**Figure S1B**), suggesting that non-specific activity of the UCP1 antibody may lead to false-positive staining in mammary glands.

**Figure 1:**
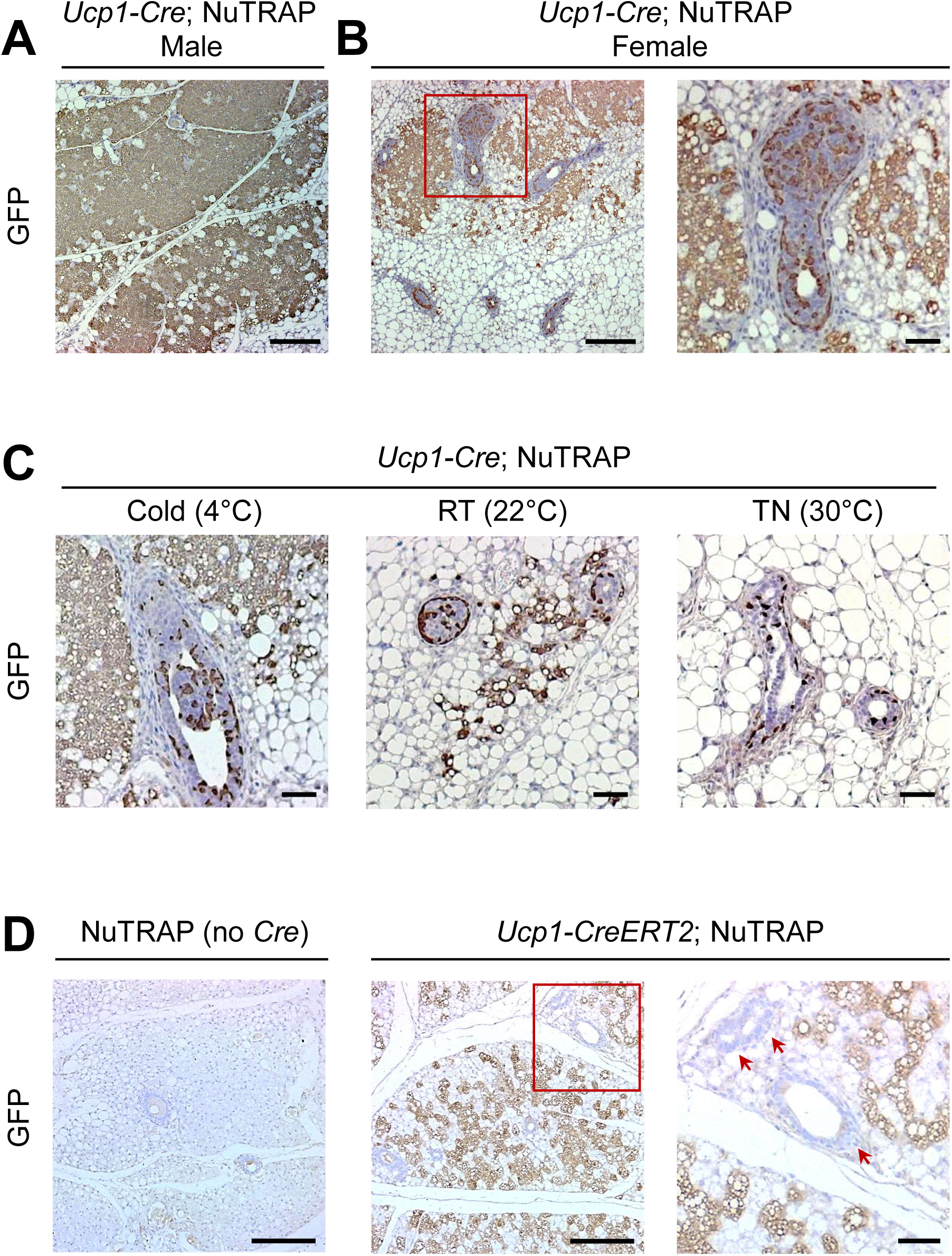
*Ucp1-Cre* expression but not endogenous *Ucp1* in the mammary glands of female mice. (A, B) Immunohistochemical staining for GFP in iWAT depots from *Ucp1-Cre*; NuTRAP male (A) and female mice (B). (C) Immunohistochemical staining for GFP in mammary glands within the iWAT depots of female *Ucp1-Cre*; NuTRAP mice under cold (4°C, 1 week), room temperature (RT; 22°C), and thermoneutrality (TN; 30°C, 4 weeks) conditions. (D) GFP Immunohistochemical staining in the iWAT of Cre-negative control (NuTRAP) and *Ucp1-CreERT2*; NuTRAP female mice after tamoxifen injection during 1-week cold exposure. Red arrows indicate mammary glands. Scale bar: 200 µm (A, B, D), 50 µm (C, high-magnification inset images in B, D).

To determine whether UCP1 is expressed in mammary glands, we employed a different approach using NuTRAP crossed with tamoxifen-inducible *Ucp1-CreERT2* mice, which allows GFP-labeling of cells with active *Ucp1* expression at the time of tamoxifen injection. We exposed female *Ucp1-CreERT2*; NuTRAP mice to cold temperatures for 1 week and administered tamoxifen for three consecutive days during cold exposure. Immunohistochemical staining using a GFP antibody revealed that GFP was expressed exclusively in beige adipocytes but not in the mammary glands within the iWAT of *Ucp1-CreERT2*; NuTRAP mice (**Figure 1D**). Immunofluorescence co-staining showed that most of beige adipocytes were positive for both UCP1 and GFP (**Figure S1C**). In contrast, mammary gland cells were only positive for UCP1, likely due to non-specific staining (**Figure S1C**). These results indicate that UCP1 is not actively expressed in the mammary glands of adult female mice. Instead, they suggest the possibility that *Ucp1-Cre* may have been once activated in mammary epithelial cell lineages during earlier development.

### 3.2. *Ucp1-Cre* expression is initiated in multiple tissues during embryonic development

To determine when *Ucp1-Cre* becomes active in the mammary glands during development, we examined GFP expression in developing *Ucp1-Cre*; NuTRAP embryos. Through gross visual inspection of developing mouse embryos, we identified GFP expression in the mammary epithelial buds on embryonic day 13.5 (E13.5) (**Figures 2A, 2B**). We found GFP expression in other areas, including the brain, eyes, whisker follicles, and internal organs (**Figure 2A, 2C**). Subsequently, we performed GFP immunochemical staining on sagittal or cross sections of the E13.5 embryos. We observed GFP expression in the epithelial layer of the choroid plexus in the forebrain (lateral ventricles) and hindbrain (fourth ventricles) (**Figure 2D**), renal tubular epithelial cells in the kidney (**Figure 2E**), scleral cells in the eye (**Figure 2F**), and the epithelium of the saccule, utricle, and semicircular ducts in the inner ear (**Figure 2G**). Interestingly, GFP expression was not yet detected in the developing BAT even at a later developmental stage of E16.5 (**Figure 2H**). UCP1 immunohistochemical staining failed to reveal any specific signals apart from high non-specific background signals throughout the entire E13.5 embryo sections (**Figure S2**). These results suggest that *Ucp1-Cre* might have been activated prior to E13.5 in various types of tissues, including the brain, kidneys, eyes, and ears.

**Figure 2:**
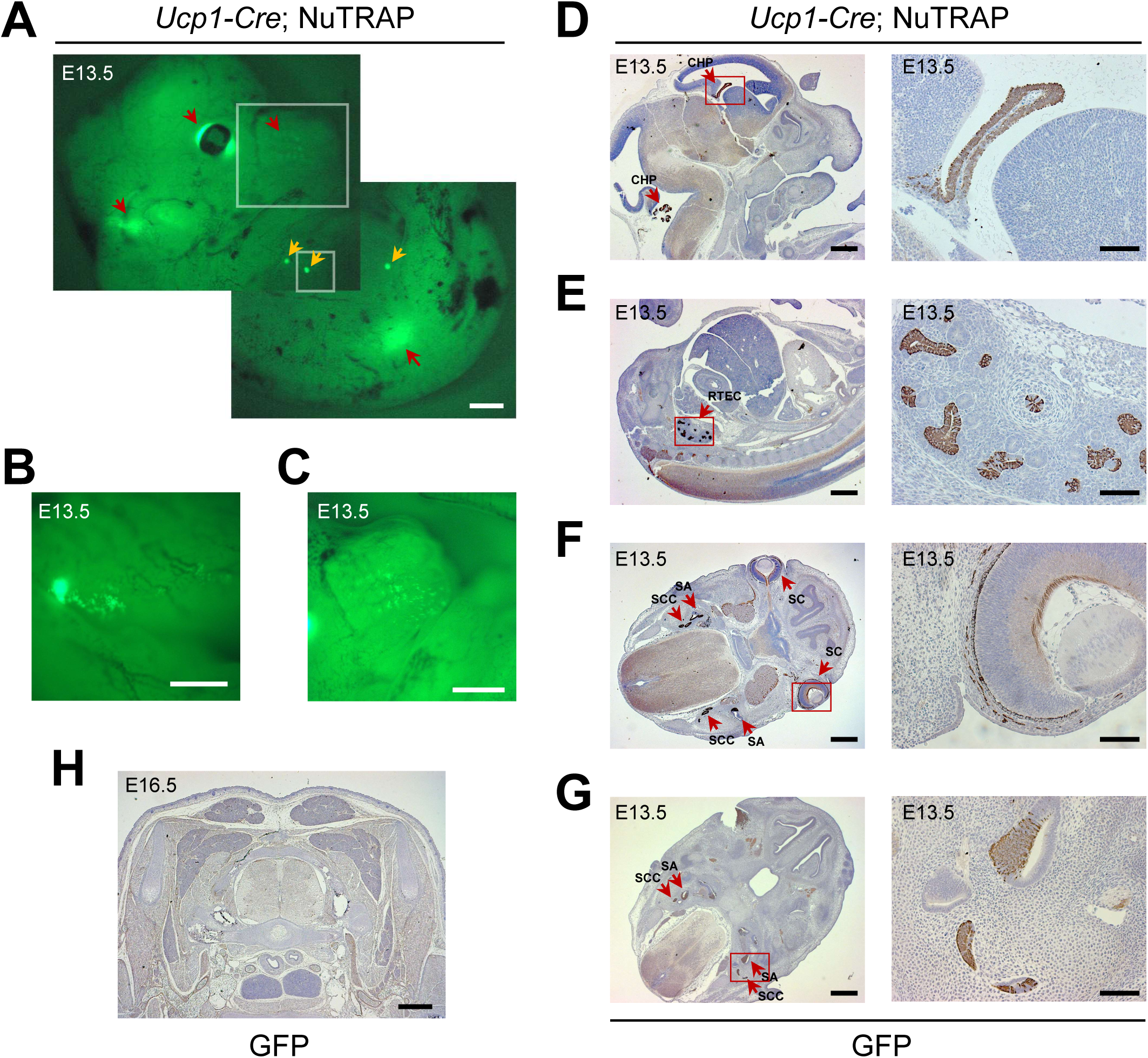
*Ucp1-Cre* activation in multiple tissues during embryonic development. (A–C) Visualization of GFP fluorescence in the intact *Ucp1-Cre*; NuTRAP embryo on embryonic day 13.5 (E13.5). Whole embryo at low magnification (A). Brain (B) and whisker follicles (C) at high magnification. Orange arrows indicate expression of GFP in the developing mammary glands. Red arrows indicate expression of GFP in the brain, eyes, whisker follicles, and internal organs. (D–G) Immunohistochemical staining for GFP in sagittal (D, E) and cross sections (F, G) of E13.5 *Ucp1-Cre*; NuTRAP embryos. (H) No GFP-positive signals in developing brown adipose tissue of E16.5 *Ucp1-Cre*; NuTRAP embryo. Scale bar: 500 µm (A, B, D–G, H), 200 µm (C), 50 µm (high-magnification inset images in D–G). CHP = choroid plexus; RETC = renal tubular epithelial cells; SC = scleral cells; SA = saccule/utricle; SCC = semi-circular canal.

To determine whether UCP1 is actively expressed in these non-adipose tissues in adult mice, we analyzed the mRNA expression of *Ucp1* and *GFP* in various tissues from the female *Ucp1-CreERT2*; NuTRAP mice exposed to cold. We examined the kidney and hypothalamus, where we and others observed *Ucp1-Cre* expression [8], along with the *Ucp1-Cre*-negative liver. As positive controls, we included BAT and iWAT with *Ucp1* expression. Both BAT and iWAT exhibited high levels of *Ucp1* expression both in Cre-negative NuTRAP mice and *Ucp1-CreERT2*; NuTRAP mice **(Figure S3A)**. *GFP* expression was strongly induced specifically in *Ucp1-CreERT2*; NuTRAP mice **(Figure S3B)**. In contrast, we detected very low levels of *Ucp1* in the kidney, hypothalamus, and liver, and *GFP* expression was not induced in these tissues from *Ucp1-CreERT2*; NuTRAP mice (**Figure S3A, S3B**). These data suggest that *Ucp1* is unlikely to be expressed in these non-adipose tissues in adult mice, and even if it is, its abundance is low and expressed only transiently.

### 3.3. sn/scRNA-seq data suggest potential endogenous UCP1 expression in mammary glands

To investigate the expression of endogenous UCP1, we utilized publicly available sn/scRNA-seq databases [25–29]. Our analysis of the snRNA-seq data obtained from adult mouse WATs demonstrated prominent *Ucp1* expression in adipocyte populations (**Figure 3A–3C**). Furthermore, we observed *Ucp1* expression, albeit in very low cell numbers, in the female epithelial cells (**Figure 3C**), which are myoepithelial cells of female mammary glands. Of note, *Ucp1* expression was also detected in other cell types, such as macrophages, male epithelial cells, and mesothelial cells (**Figure 3C**). We subsequently extended our investigation to human tissues by analyzing scRNA-seq data available from the Human Protein Atlas [20]. Consistent with the mouse data, *UCP1* expression was detected in breast glandular and myoepithelial cells in humans (**Figure 3D**). These findings suggest the possibility of endogenous UCP1 expression in mammary glands in adults.

**Figure 3:**
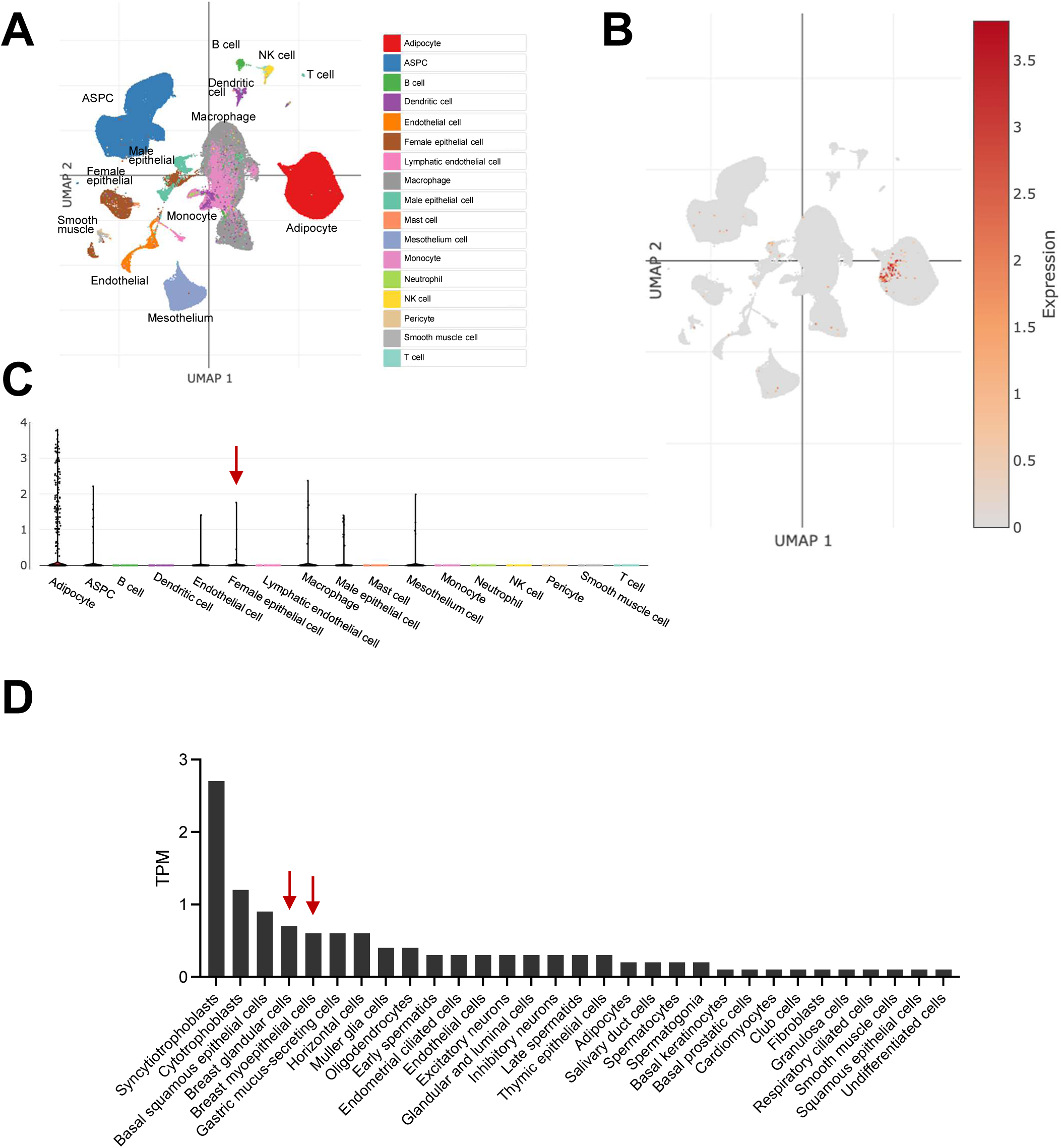
Single nucleus/cell RNA-seq analysis of UCP1 expression in mouse and human mammary epithelial cells. (A) Uniform manifold approximation and projection (UMAP) projection of cell clusters from mouse WATs. (B) Feature plot showing *Ucp1*-expressing cells. (C) Violin plot showing *Ucp1* expression across different cell clusters in the mouse WAT dataset. (D) *UCP1* expression in various cell types from human scRNA-seq data.

### 3.4. UCP1 deficiency has no impact on mammary gland structure or functions

Given the potential expression of UCP1 in myoepithelial and luminal cells in the mammary glands, we sought to investigate whether UCP1 is involved in mammary gland development and function. We first examined the structure of mammary glands in female UCP1-deficient mice, both non-pregnant and lactating post-delivery. In both cases, *Ucp1*-KO female mice exhibited abundant multilocular beige adipocytes in iWAT (**Figure 4A**), as previously reported [23,24]. However, the structure of the mammary glands, including the morphology of ductal and alveolar regions, appeared similar between *Ucp1*-KO and control heterozygous mice, regardless of whether they were in a non-pregnant or lactating state (**Figure 4A**). To assess the functionality of the mammary glands, we crossed *Ucp1*-KO or heterozygous female mice with wild-type male mice and monitored the body weight of the resulting pups during a two-week lactation period. *Ucp1* heterozygous female mice served as better controls than wild-type female mice when crossed with wild-type male mice since 50% of their pups shared the same genotypes as those from *Ucp1*-KO mothers. This approach minimized the effects of genotype differences in the pups. There was no significant difference in the body weights of the pups between the different maternal genotypes (**Figure 4B**), indicating a similar nutritional content from lactation and comparable functionality of the mammary glands between genotypes. Taken together, these results suggest that the loss of UCP1 may not have a significant impact on mammary gland development and function.

**Figure 4:**
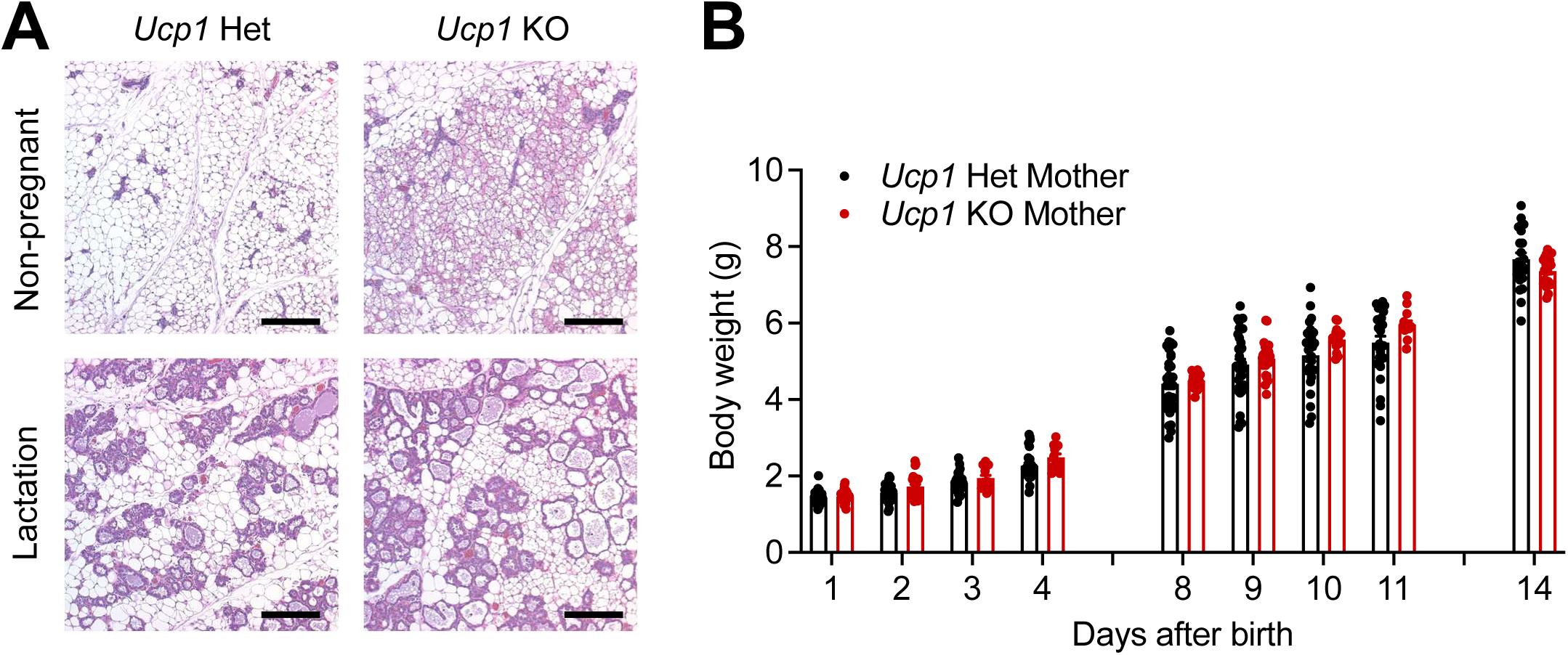
Normal development and function mammary glands in *Ucp1*-deficient mouse. (A) H&E staining of iWAT sections from *Ucp1* heterozygous (Het) and KO mice under non-pregnant (top) and lactation states (bottom). Scale Bar: 200 µm. (B) Body weights of pups born to lactating female mice with *Ucp1* Het (*n* = 27) or KO (*n* = 18) for 2 weeks after birth. These female mice were mated with wild-type male mice.

## 4. DISCUSSION

UCP1 is widely recognized as one of the marker genes exclusively expressed in brown and beige adipocytes. *Ucp1-Cre* mice have been a crucial tool for studying thermogenic adipocytes, particularly for generating a variety of mouse models with specific genetic perturbations in brown and beige adipocytes. However, the utility of *Ucp1-Cre* mice has recently been questioned due to the finding of Cre activity in additional tissues, including the kidneys, adrenal glands, and hypothalamus, all of which contribute to energy homeostasis [8]. Therefore, it is important to carefully investigate *Ucp1-Cre* expression and exercise caution when interpreting findings from studies using this model.

In this study, using *Ucp1-Cre*-driven NuTRAP reporter mice, we found that *Ucp1-Cre* is expressed in myoepithelial and luminal cell layers within the mammary glands of female mice. This suggests the possibility of endogenous UCP1 expression in cell types other than adipocytes within adipose tissues. This expression in the mammary gland remained unaffected by changes in ambient temperature, unlike the cold-inducibility of *Ucp1* in adipocytes. Our initial UCP1 immunostaining analysis indicated the presence of UCP1 in mammary glands. However, the subsequent experiments using *Ucp1*-KO mice suggested that UCP1 staining in mammary glands was likely a false positive due to non-specific immunoreactivity of the UCP1 antibody. Furthermore, the failure to induce GFP expression in the mammary glands of tamoxifen-inducible *Ucp1-CreERT2*; NuTRAP mice supported the lack of active UCP1 expression. These results prompted us to investigate the timing of *Ucp1-Cre* expression during development. We found that *Ucp1-Cre* becomes active in the developing mammary buds by E13.5. These findings suggest that UCP1 expression may be an early developmental event. Interestingly, when examining sc/snRNA-seq data from both adult mouse and human tissues, we found low-level *UCP1* expression in a small number of cells from the mammary glands. These results suggest the possibility that UCP1 may be transiently expressed at low levels in adult mammary glands. However, it is also plausible that *Ucp1*-expressing mammary gland cells in the sc/snRNA-seq data represent doublets between *Ucp1*-positive adipocytes and *Ucp1*-negative mammary gland cells. Given that *Ucp1* was similarly detected in other cell types, such as macrophages, male epithelial cells, and mesothelial cells, these data do not provide definitive conclusions regarding endogenous UCP1 expression. In addition, *Ucp1*-KO mice exhibited normal mammary gland development and functionality. As such, it remains an unanswered question whether UCP1 is expressed in and functionally important for adult mammary glands.

Previous studies documented the existence of a specific type of adipocyte known as pink adipocytes, which were proposed to interconvert between mammary gland epithelial cells and subcutaneous adipocytes [30,31]. However, a recent cell tracing study has challenged this notion and provided evidence against adipocyte-to-epithelial cell conversion [32]. Nonetheless, it is possible that UCP1-expressing epithelial cells within the mammary gland are related to the proposed pink adipocytes. In this scenario, rather than pink adipocytes capable of converting into epithelial cells, these UCP1-expressing cells likely represent a committed cell type akin to myoepithelial and ductal cells. Although no functional defects were observed in the mammary glands of *Ucp1*-KO mice, it is possible that UCP1 may serve specific functions under certain conditions, such as warming milk within the mammary ducts during low temperatures. Further investigations are required to explore this possibility.

A recent study by Patel et al. discovered that mammary luminal epithelial cells produce paracrine secretory factors, named mammokines, such as lipocalin 2, which inhibit UCP1 expression in adipocytes within the iWAT of female mice [33]. This study raises some intriguing questions. Are the mammary epithelial cells expressing *Ucp1-Cre* the same as or functioning with those that secrete mammokines? Could mammokines switch off UCP1 expression in mammary ducts by acting as autocrine factors? Such a mechanism might explain the lack of UCP1 expression in these cells in adult female mice. Further investigation is warranted to determine whether UCP1 expression in the mammary gland during its development plays a role in iWAT homeostasis in female mice.

We observed *Ucp1-Cre* expression in additional tissues that have not been previously reported [8], including the eyes, ears, and whisker follicles. This finding suggests that the *Ucp1-Cre* expression pattern is broader than previously believed. Interestingly, we noticed that *Ucp1-Cre* is consistently expressed in epithelial cell layers within these tissues. This observation implies that UCP1 may have functional implications in this particular cell type, which warrants further investigation in future studies. However, we cannot completely exclude the possibility that the expression of *Ucp1-Cre* in epithelial cells could be an artifact resulting from the bacterial artificial chromosome-derived construct and its insertion in the genome. Using *Ucp1-CreERT2;* NuTRAP mice, we found that *Ucp1* is not actively expressed in these non-adipose tissues, similar to mammary epithelial cells, suggesting that the role of UCP1 in these tissues may be related to developmental processes. It is important to note that while *Ucp1-Cre* is broadly expressed, *Ucp1-CreERT2* induces Cre recombination specifically in adipocytes upon tamoxifen administration. None of these non-adipose tissues or mammary epithelial cells that we tested displayed significant Cre recombination. This finding is contrary to the previous report suggesting active *Ucp1* expression in the hypothalamus of the adult mouse brain [8]. The discrepancy may be due to the different methodologies, specifically the use of tamoxifen-inducible genetic reporters versus Cre-inducible viral reporter injection. Regardless, our results demonstrate that *Ucp1-CreERT2* can serve as a useful tool for circumventing complications arising from the activation of the *Ucp1* promoter in non-adipose tissues during embryonic development.

In conclusion, our findings reveal that the expression pattern of UCP1 is wider and more dynamic than previously understood. Consequently, it is essential to exercise caution when interpreting data and designing experiments using *Ucp1-Cre* mice models. Furthermore, the utilization of additional complementary tools, such as the tamoxifen-inducible *Ucp1-CreERT2* system, should receive thorough consideration during experimental design.

## AUTHOR CONTRIBUTIONS

H.C.R. conceived and designed the study. H.C.R., K.K., J.W., and H.G.K. performed experiments. H.C.R., K.K., and E.D.R. interpreted the data. H.C.R. and J.S. performed computational analysis. H.C.R. and K.K. wrote the manuscript with inputs from all the other authors.

## ACKNOWLEDGEMENTS

We thank Christian Wolfrum for providing *Ucp1-CreERT2* mice. We are grateful for the support from the Histology Core at Beth Israel Deaconess Medical Center and Indiana University School of Medicine.

## Abbreviations

WAT: White adipose tissue
BAT: Brown adipose tissue
UCP1: Uncoupling protein 1
iWAT: Inguinal white adipose tissue
KO: Knockout
sc/snRNA-seq: Single-cell/nucleus RNA-seq
NuTRAP: Nuclear tagging and Translating Ribosome Affinity Purification
E13.5: Embryonic day 13.5
E16.5: Embryonic day 16.5

