## Supplementary Figures for "Uncoupling protein 1-driven Cre (*Ucp1-Cre*) is expressed in the epithelial cells of mammary glands and various non-adipose tissues"

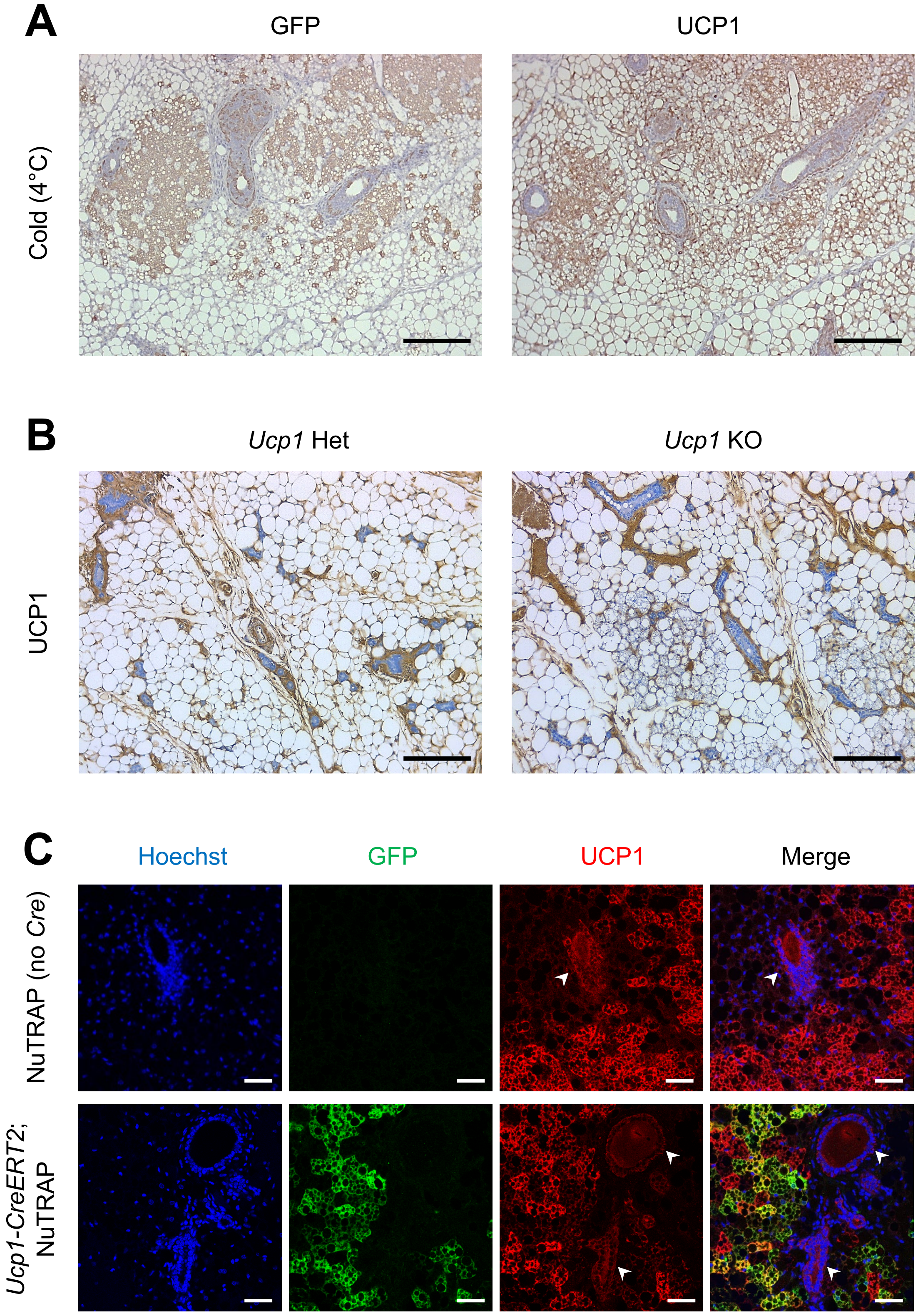
A
GFP
UCP1
Cold (4°C)
B
Ucp1 Het
Ucp1 KO
UCP1
C
Hoechst
GFP
UCP1
Merge
NuTRAP (no Cre)
Ucp1-CreERT2;
NuTRAP

**Figure S1: Non-specific staining of UCP1 antibody in iWAT.** (A) Immunohistochemical staining for GFP and UCP1 in consecutive iWAT sections from a female *Ucp1-Cre*; NuTRAP mouse exposed to cold (4°C) for 1 week. (B) Immunohistochemical staining using UCP1 antibody (Abcam, ab10983) in the iWAT of *Ucp1* heterozygous (Het) and knockout (KO) female mice. (C) Confocal images of double immunofluorescence staining for GFP and UCP1 in the iWAT from Cre-negative control (NuTRAP) and *Ucp1-CreERT2*; NuTRAP female mice after tamoxifen injection during 1-week cold exposure. White arrowheads indicate UCP1-positive mammary glands. Scale bar: 200 μm (A, B), 50 μm (C).

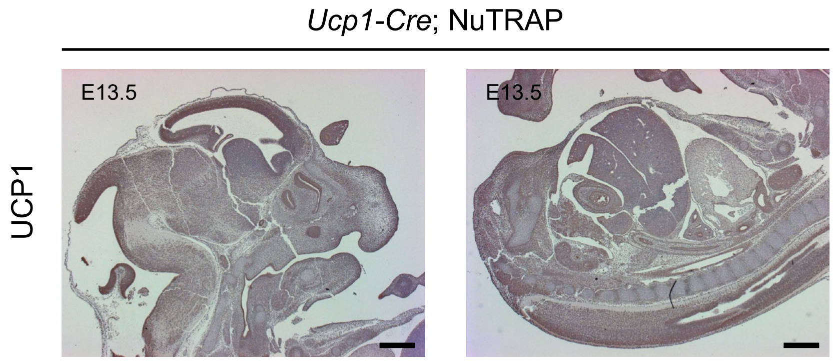


**Figure S2: High background staining of UCP1 antibody in developing embryos.** Immunohistochemical staining for UCP1 in sagittal sections of the upper (left) and lower (right) body of developing embryos of *Ucp1-Cre*; NuTRAP on embryonic day 13.5 (E13.5). Scale bar: 500 µm.

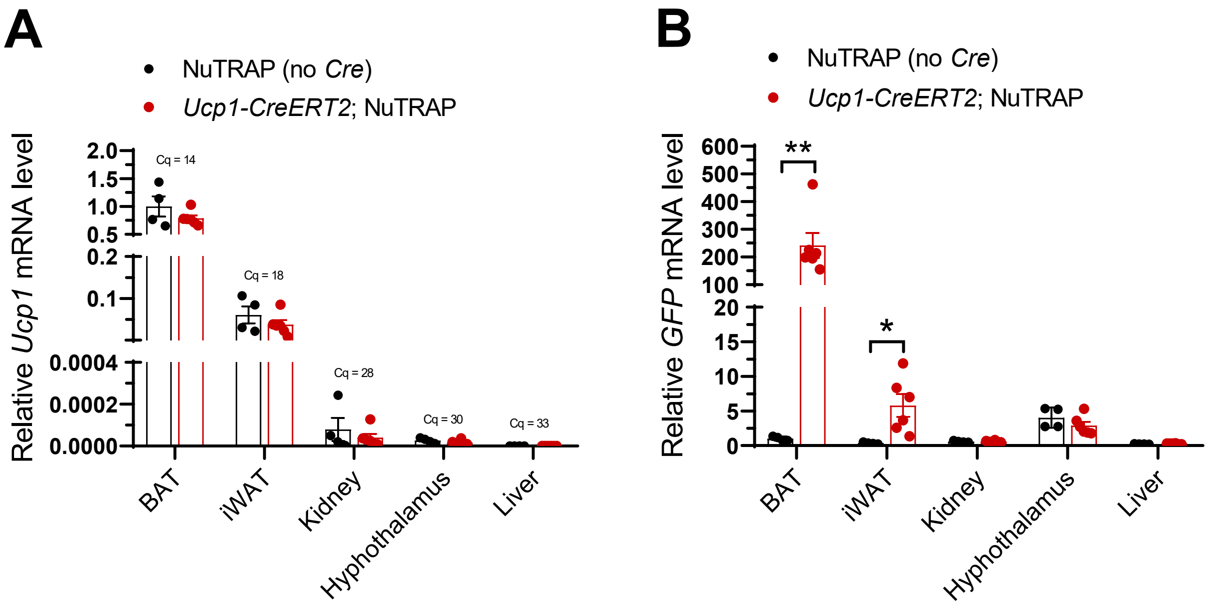


**Figure S3: Expression of *Ucp1* and *GFP* mRNA in various tissues of *Ucp1-CreERT2*; NuTRAP female mice exposed to cold.** Relative levels of *Ucp1* (A) and *GFP* (B) mRNA expression in the BAT, iWAT, kidney, hypothalamus, and liver from Cre-negative control (NuTRAP; n = 4) and *Ucp1-CreERT2*; NuTRAP female mice (n = 6) after tamoxifen injection during 1-week cold exposure. The quantification of cycle (Cq) values from quantitative real-time PCR for *Ucp1* mRNA are presented for each tissue. Data are shown as mean ± SEM, 2-tailed unpaired Student's *t* test. *$P < 0.05$, **$P < 0.01$.
